# Mitochondrial Fusion Suppresses Pancreatic Cancer Growth via Reduced Oxidative Metabolism

**DOI:** 10.1101/279745

**Authors:** Meifang Yu, Yanqing Huang, Amit Deorukhkar, Tara N. Fujimoto, Suman Govindaraju, Jessica M. Molkentine, Daniel Lin, Ya’an Kang, Eugene J. Koay, Jason B. Fleming, Sonal Gupta, Anirban Maitra, Cullen M. Taniguchi

## Abstract

Pancreatic cancer is a highly lethal disease whose aggressive biology that is driven by mitochondrial oxidative metabolism. Mitochondria normally form a network of fused organelles, but we find that patient-derived and genetically engineered murine pancreatic cancer cells exhibit highly fragmented mitochondria with robust oxygen consumption rates (OCR). When mitochondrial fusion was activated by the genetic or pharmacological inhibition Drp1, the morphology and metabolism of human and murine pancreatic cancer cells more closely resembled that of normal pancreatic epithelial cells. This reduced metabolism was correlated with slower tumor growth, fewer metastases, and enhanced survival in a syngeneic orthotopic model. Similarly, directly activating mitochondrial fusion by overexpression of Mfn2 also reduced tumor growth and metastases. Mitochondrial fusion in pancreatic cancer cells was associated with reduced mitochondrial mass and Complex I expression and function. Thus, these data suggest that enhancing mitochondrial fusion through Drp1 inhibition or enhanced Mfn2 expression or function has strong tumor suppressive activity against pancreatic cancer and may thus represent a highly novel and efficacious therapeutic target.

## Introduction

Patients with pancreatic ductal adenocarcinoma (PDAC) have a 5-year survival of less than 7% (Kleeff et al., 2016), in part because this form of cancer is highly resistant to nearly all treatments and metastasizes frequently. Recent work has shown that pancreatic cancer is metabolically robust and depends on oxidative phosphorylation for growth and progression (Sancho et al., 2015; Viale et al., 2014). Furthermore, treatment-resistant clones are likely to have increased mitochondrial mass and respiration rates (Viale et al., 2014).Furthermore, PDAC may also fuel its dependence on the mitochondria by usurping nearby stromal cells to fuel the mitochondrial tricarboxylic acid (TCA) cycle (Sousa et al., 2016).

Mitochondria are not static organelles and constantly change their morphology in response to stress and energetic needs of the cell (Vyas et al., 2016). In most normal tissues (Luchsinger et al., 2016) and some tumors, mitochondria are fused together to optimize oxidative phosphorylation and to facilitate repair of damaged mtDNA. Cell that are transformed by H-ras, however, exhibit enhanced mitochondrial fission (Kashatus et al., 2015) through a process dependent on MEK, which phosphorylates the key regulator dynamin-related protein1 (Drp1). Despite these higher levels of fragmented mitochondria, Ras-transformed cancer cells exhibited enhanced growth and metastasis. Whether mitochondrial fission supports higher metabolic rates in pancreatic cancer is unknown, since this process is normally associated with lower metabolic rates.

To resolve this apparent paradox, we studied the mitochondrial dynamics and metabolism from primary patient tumors and murine pancreatic cancer lines and found that aggressive pancreatic cancers exhibit high levels of mitochondrial fission and oxidative phosphorylation. Conversely, we found that a cell line from a more indolent cancer exhibited more mitochondrial fusion. When we enhanced mitochondrial fusion by direct antagonism of Drp1 or by expressing mitofusin-1 (Mfn2), pancreatic cancer growth and metastasis was suppressed. Thus, promoting mitochondrial fusion may be a promising strategy for treating pancreatic cancer.

## Results

### Mitochondrial fission is associated with higher oxidative metabolism in pancreatic cancer

We analyzed mitochondrial dynamics in six primary pancreatic lines (Kang et al., 2015) to determine whether fission, fusion or an intermediate phenotype was present. We found that normal human pancreatic normal epithelial (HPNE) cells had elongated fused mitochondria as determined by mitotracker staining and confocal microscopy (Figure 1A and quantitated in Figure 1B). A similar analysis in other primary cell lines established directly from resected patient tumors at our institution revealed that pancreatic tumors nearly always exhibited high levels of mitochondrial fission (Figure 1A and quantitated in Figure 1B).There was only one cell line, MDA-PATC69 (PATC69), that did not exhibit a florid fission phenotype and instead exhibited a modest amount of mitochondrial fusion. We confirmed the morphological observations of mitochondria by electron microscopy in HPNE, PATC118 and PATC69 cells (Figure 1C).

**Figure 1.**
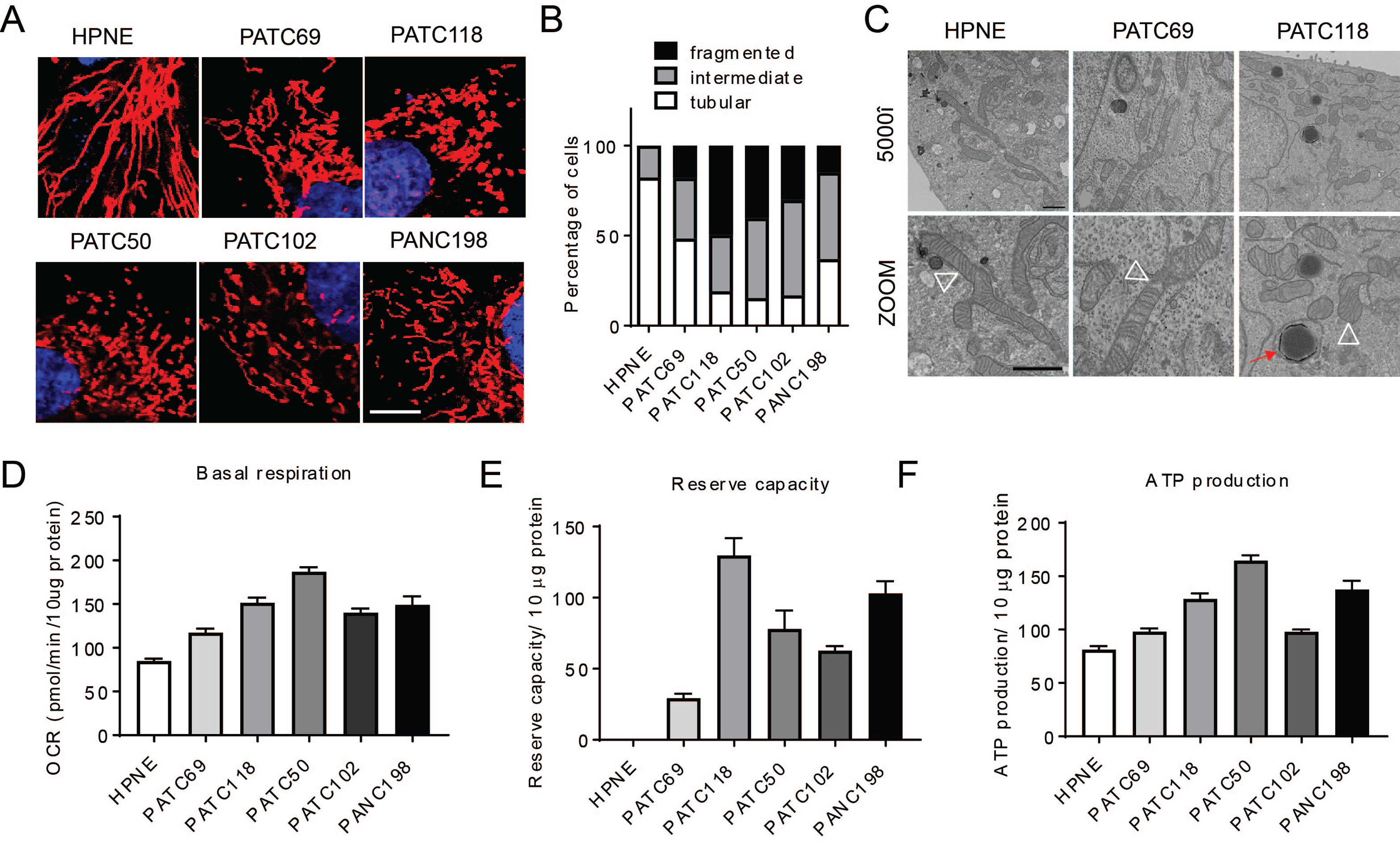
Mitochondrial fission is associated with higher oxidative metabolism in pancreatic cancer. Patient-derived pancreatic cells and human pancreas normal epithelial were cultured for mito stress assay and mitochondrial morphology measurement. (A) Mitochondria staining of normal pancreas epithelial cells (HPNE) and five different patient– derived pancreatic cancer lines show most cell lines utilize mitochondrial fission, with morphology quantified in (B) (n=100-200 cells per line). Scale bar=10µm. (C) Transmission electron microscopy (TEM) image of mitochondrial morphology. Pictures were taken at 5000× and zoomed in for detail. White open triangle points to mitochondrion, scale bar=1µm. (D) Mita stress assay of these pancreatic cancer cells. Quantification of metabolic parameters from mito stress assay of (D) Basal oxygen consumption rate, (E) reserve capacity, and (F) ATP production. Data were normalized to protein concentration. Statistic analyses are presented in Figure S1. Data are presented as mean± SEM. *p<0.05, **p<0.01, ***p<0.001, ****p<0.0001.

To understand whether mitochondrial fission could support higher levels of oxidative metabolism, we measured oxygen consumption rates on several of the cell lines using Seahorse extracellular flux analyzer. We found that the cells lines from patients with a more aggressive clinical course had higher basal OCR (Figure 1D), higher spare respiration capacity (Figure 1E), and more ATP production than HPNE or the more indolent PATC69, and these comparisons all reached statistical significance (Figure 1F, with complete *p-*values in Figure S1C, 1D and 1E). The extracellular acidification rate (ECAR), which serves as a proxy for glycolysis, was not significantly different between these lines (Figure S1A, B). Interestingly, this patient that donated the PATC69 cell line with a mitochondrial fusion phenotype had a more indolent course of cancer with an overall survival (OS) of 33.3 months, which exceeded the lifespan of the other patient donors, who unfortunately only survived between 4-16 months (Figure S1F).

We further confirmed the fission and fusion phenotypes with immunoblot analysis of key effectors of mitochondrial fission and fusion (Detmer and Chan, 2007). Most notably, we find that Drp1 expression and phosphorylation are increased in the PATC118 and Mfn2 expression is decreased compared to PATC69 and normal HPNE controls (Figure 2A and quantified in Figure 2B).These molecular changes in Drp1 and Mfn2 are concordant with the high levels of fragmented mitochondria (Figure 1A and 1C). We also saw increased OPA1 expression in cancer cells compared to normal HPNE control, did not observe any change in other key molecules such as Mfn1 (Figure S2A, 2B).

**Figure 2.**
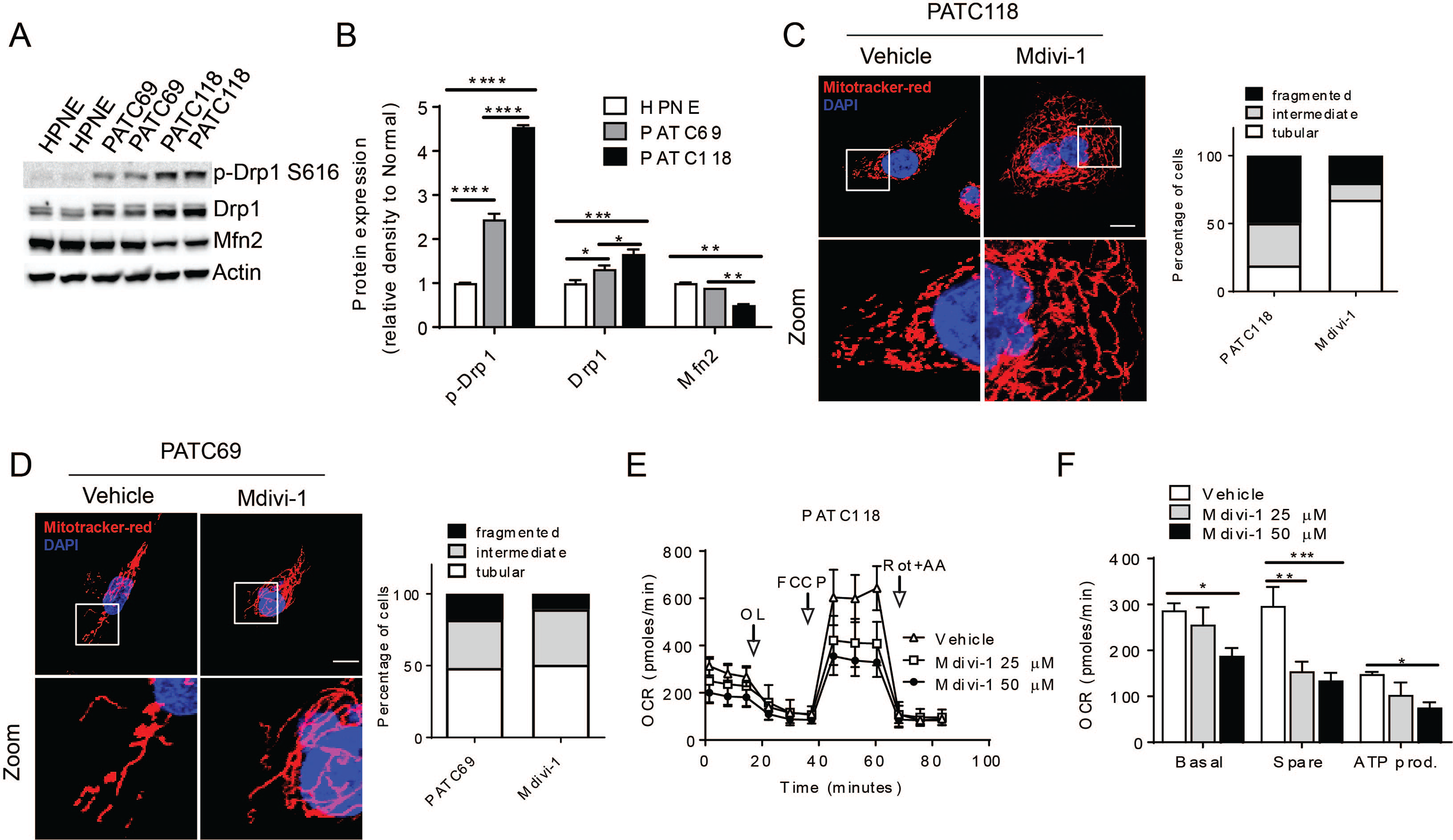
Induction of mitochondrial fusion decreases oxygen consumption in primary pancreatic cell lines. (A) lmmunoblot analysis of mitochondrial dynamics related proteins in normal line and pancreatic cancer lines. Each lane represents a representative biological replicate from a separate experiment. Densitometric quantification is shown in (B), with an n= 3 for each immunoblot. Mitochondrial morphology of (C) PATC118 cells and (D) PATC69 after treatment with Mdivi-1 or vehicle control. Quantification of mitochondrial morphology is shown in insert panels, n=100-200 cells were counted per line. Scale bar= 10µm. (E) PATC118 cells showed decreased mitochondrial respiration after treatment with 25 or 50µM Drp1 inhibitor, and parameters of the mito stress test are quantified in (F). Data were normalized to protein concentration. See also Figure S2. Data are presented as mean± SEM. *p<0.05,**p<0.01,***p<0.001, ****p<0.0001.

### Induction of mitochondrial fusion decreases oxygen consumption in primary pancreatic cell lines

Mdivi-1 is a known pharmacologic inhibitor of Drp1 function (Cassidy-Stone et al., 2008), and to understand if this drug could alter mitochondrial dynamics and metabnolism in pancreatic cancer, we treated PATC118 (Figure 2C) and PATC69 (Figure 2D) for 24 hours. We found that Mdivi-1 normalized the morphology of PATC118 cells to a more elongated form, whereas there was no effect in the already elongated mitochondria of PATC69 cells. Furthermore, we observed that this enhanced mitochondrial fusion in PATC118 decreased basal OCR, spare consumption, and total ATP production in a dose-dependent fashion (Figure 2E and quantified in 2F). Conversely, there were no oxygen consumption changes in PATC69 cells (Figure S2C, 2D), suggesting that alteration in fission and fusion, and not merely Drp1 inhibition was required for changes in metabolism.

### Genetic or pharmacological induction of mitochondrial fusion suppresses pancreatic cancer growth and improves survival

To understand how mitochondrial dynamics could alter the biology of pancreatic cancer, we turned to a murine model using KPC cells (*Kras*^LSL/+^; *Trp53*^R172H/+^; Pdx1^Cre/+^) syngeneic to C57BL/6 to allow more facile manipulation and in vivo interrogation. These KPC tumors grow rapidly and have a high metastatic potential (Guerra and Barbacid, 2013), and we hypothesized that they would share mitochondrial characteristics from our PATC118 line. We CRISPR-Cas9 methodology to ablate Drp1 expression, which was confirmed by Western blot (Figure 3A, for other clones, see Supplemental Figure 4). Confocal microscopy with mitotracker staining revealed that KPC cells have highly fragmented morphology (Figure 3B), but loss of Drp1 reverted more than 80% of the cells from fragmented to elongated mitochondria (Figure 3B, quantified in Figure 3C). We confirmed these morphological findings by electron microscopy (Figure S3A).

**Figure 3.**
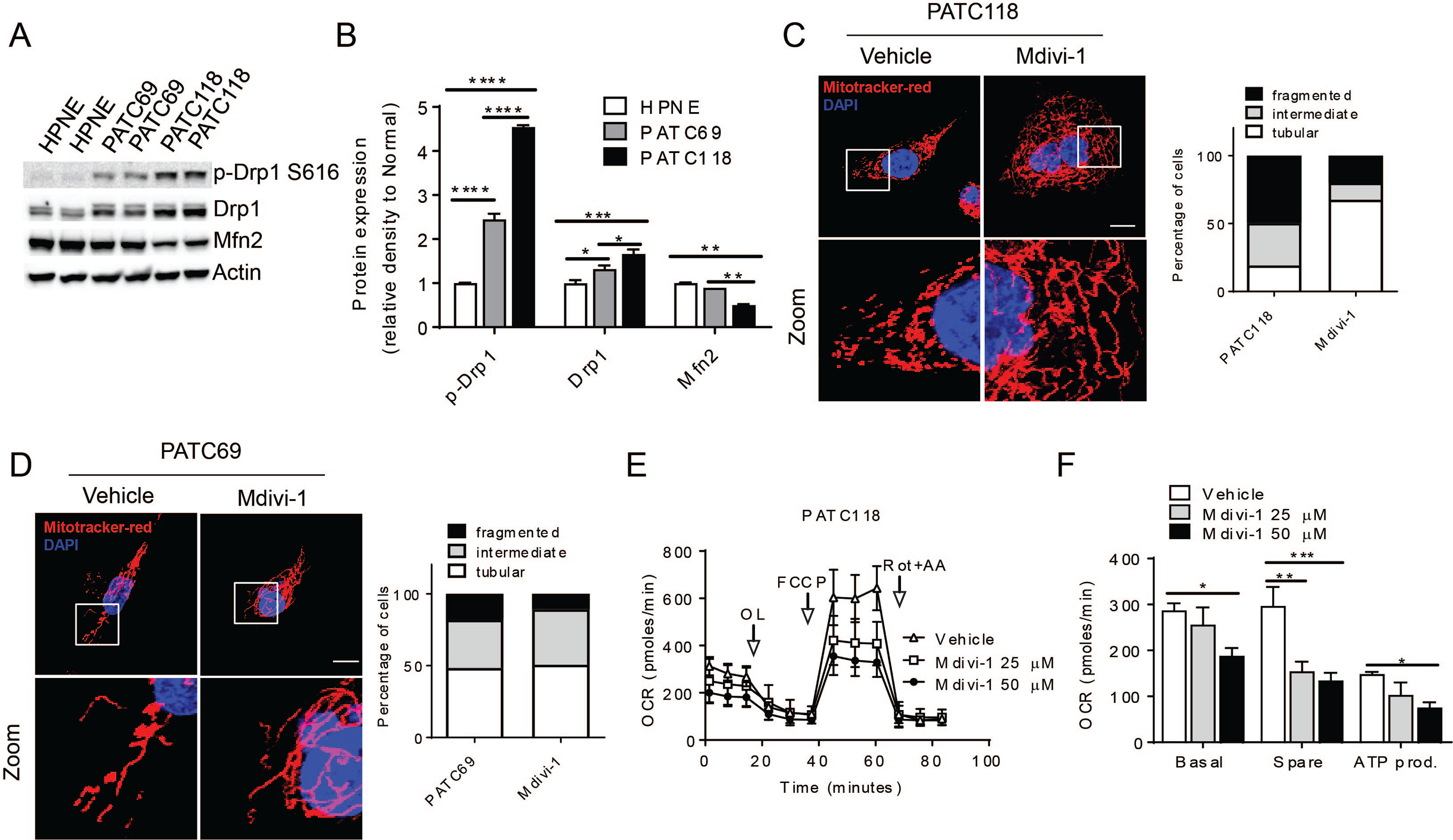
Genetic or pharmacological induction of mitochondrial fusion suppresses pancreatic cancer growth and improves survival. (A) Confirmation of Drp1 knockout by using CRISPR/Cas9 system using Western blot. (B) Loss of Drp1 (sgDRP1) induces mitochondrial fusion compared to sgGFP control, with morphoplogy quantified in (C), n=100-200 cells were counted per line. (D) Mita stress assay shows decreased basal respiration, spare respiration and ATP production in sgDrp1 cells. (E) Kaplan-Meier survival curves of C57BL/6J mice with orthotopically implanted syngeneic tumors, n=5 per cohort. (Log Rank p=0.0074).! (F) Mitochondrial morphology of KPC cells treated with Mdivi-1 and quantified in (G). Scale bar 10µm. (H) Mita stress assay of KPC treat with Mdivi-1. Data were normalized to protein concentration. (I) Tumor volume of KPC xenograft treated with Vehicle or 10mg/kg Mdivi-1, n=10 per cohort. See also Figure S3, S4. Data are presented as mean± SEM. *p<0.05, **p<0.01, ****p<0.0001.!

**Figure 4.**
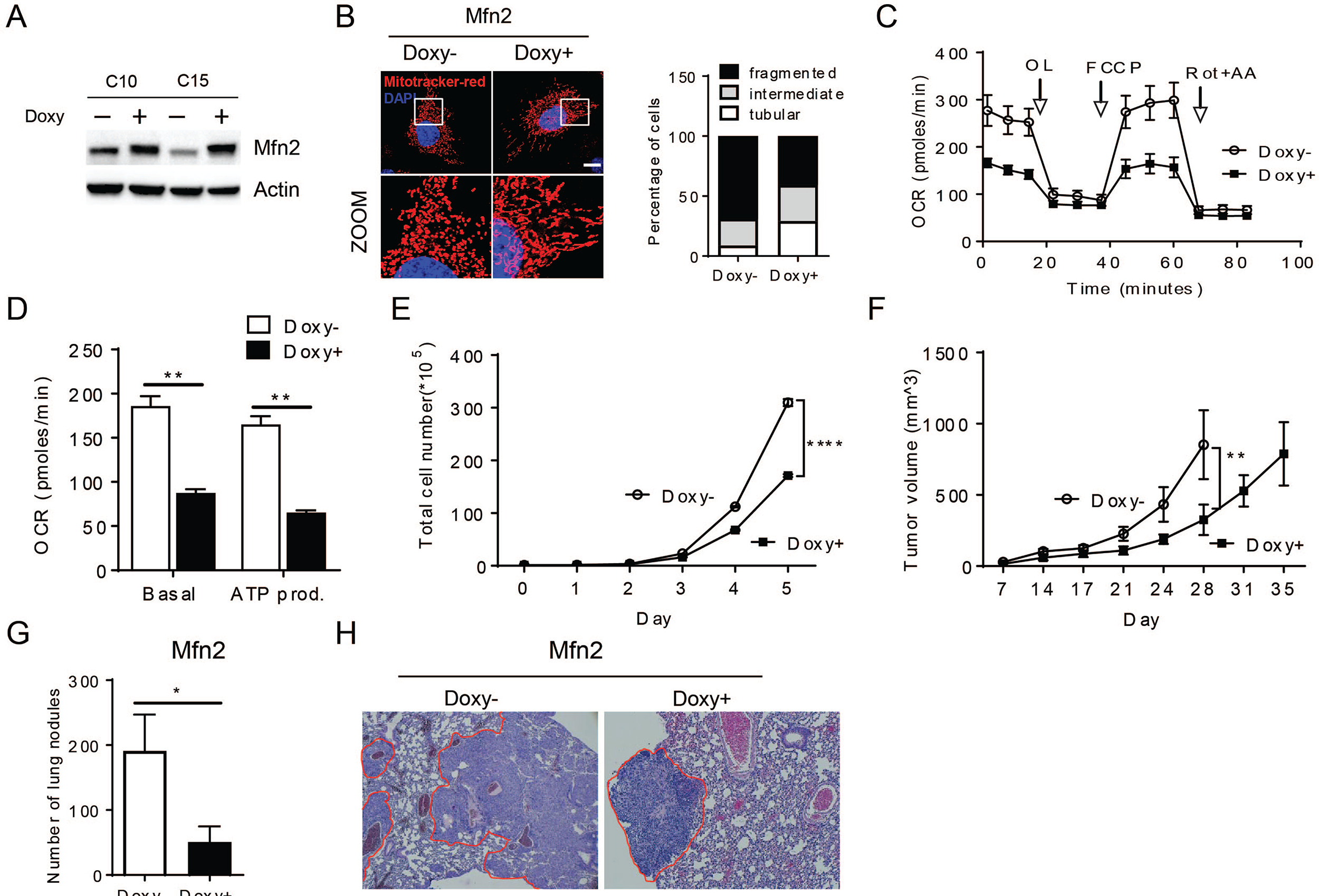
Direct induction of mitochondrial fusion with suppresses pancreatic growth and metastasis. KPC pancreatic cancer cells syngeneic to C57BL/6 with a doxycycline-inducible expression of Mfn2 were created, and expression verified by Western blot in (A). (B) Mitochondrial morphology of control group (Doxy-) and mfn2 overexpression (Doxy+), and quantified in associated panel, n= 100-200 cells were counted per line. Scale bar=10µm. (C) Mita stress assay of control group (open circle) and Mfn2-overexpressing group (closed square) and individual parameters quantified in (D). Data were normalized to protein concentration. (E) In vitro proliferation of control (open circle) and Mfn2 overexpression cells (closed square). (F) Orthotopic tumor volume in C57BL/6J mice. Tumor volume were observed twice a week after injection and represented as mean values for each group, n=5 per cohort. (G) Lung metastatic nodules found in control group (white bar) and experimental group (black bar). (H) H&E staining image (20 ×) of the lungs with metastatic nodules. See also Figure S5 Data are presented as mean± SEM. *p<0.05, **p<0.01, ****p<0.0001.A

To determine if this knockout had any functional differences in biology, we measured oxygen consumption of these sgDrp1 Cells, and found that Drp1 knockout decreased basal respiration, spare respiration capacity and ATP production compared to KPC controls (Figure 3D and S3B). This metabolic impairment, correlated with decreased *in vitro* proliferation (Figure S3C). We then performed orthotopic transplantations the Drp1 knockout cells in recipient C57BL/6 mice and found that the *in vivo* growth was also impaired (Figure S3D), which improved survival compared to KPC controls (Figure 3E). We got another set of clones of Drp1 knockout, and all the data were kept in Figure S4.

We complemented this genetic approach of inducing mitochondrial fusion with pharmacologic inhibition by Mdivi-1. We found that the addition of Mdivi1 induced mitochondrial fusion (Figure 3F and quantified in 3G). Furthermore, Mdivi-1 decreased basal oxidative metabolism, spare respiration, and ATP production in a dose-dependent manner (Figure 3H and S3E). These metabolic changes corresponded to a dose-dependent decrease *in vitro* cell growth (Figure S3F) and in vivo pancreatic cancer growth using a syngeneic flank model (Figure 3I).

### Direct induction of mitochondrial fusion suppresss pancreatic growth and metastasis

Although inhibiting mitochondrial fission through CRISPR/Cas9 or Mdivi-1 effectively induced mitochondrial fusion and suppressed growth, we wanted to confirm that the phenotype was due to the induction of fusion and not some pleiotropic effect of inhibiting Drp1. To answer this question, we created a doxycycline-inducible Mfn2 system to overexpress this gene in KPC cells. Since some previous studies had suggested that doxycycline might itself impair mitochondrial function (Moullan et al., 2015), we confirmed that doxycycline alone did not increase KPC tumor growth (Supplemental Figure 6A and 6B).However, out of an abundance of caution, we used the lowest possible concentration of doxycycline to induce effects, which was 0.15mcg/mL in the drinking water of animals. Even at this low concentration, we found robust expression of Mfn2 (Figure 4A).

The induction of Mfn2 by doxycycline produced more fused and elongated mitochondria (Figure 4B), which was correlated with lower basal OCR (Figure 4C) and ATP production (Figure 4C and quantified in 4D) compared to controls. This reduction in oxidative metabolism by Mfn2 overexpression correlated with decreased growth in vitro (Figure 4E) and in vivo in an orthotropic model (Figure 4F). Furthermore in a tail vein model of metastasis, the induction of Mfn2 significantly decreased the number of macroscopic nodules in the lung (Quantified in Figure 4G), as well as size of metastases as depicted in H&E staining (Figure 4H). These results were similar with another clone of Mfn2 overexpressing line (Figure S5A-E).

### Mitochondrial fusion decreases mtDNA and attenuate mitochondrial oxidative phosphorylation

We wondered if mitochondrial fusion induced a change in mitochondrial mass. Indeed, in human primary lines, HPNE and PATC69 displayed significantly less mitochondrial mass compared to the aggressive and hypermetabolic PATC118 line (Figure 5A). A similar analysis was performed in the multiple murine KPC lines with and without Drp1 knockout showed a decrease total mitochondrial mass as detected the mtDNA copy number(Figure 5B).

**Figure 5.**
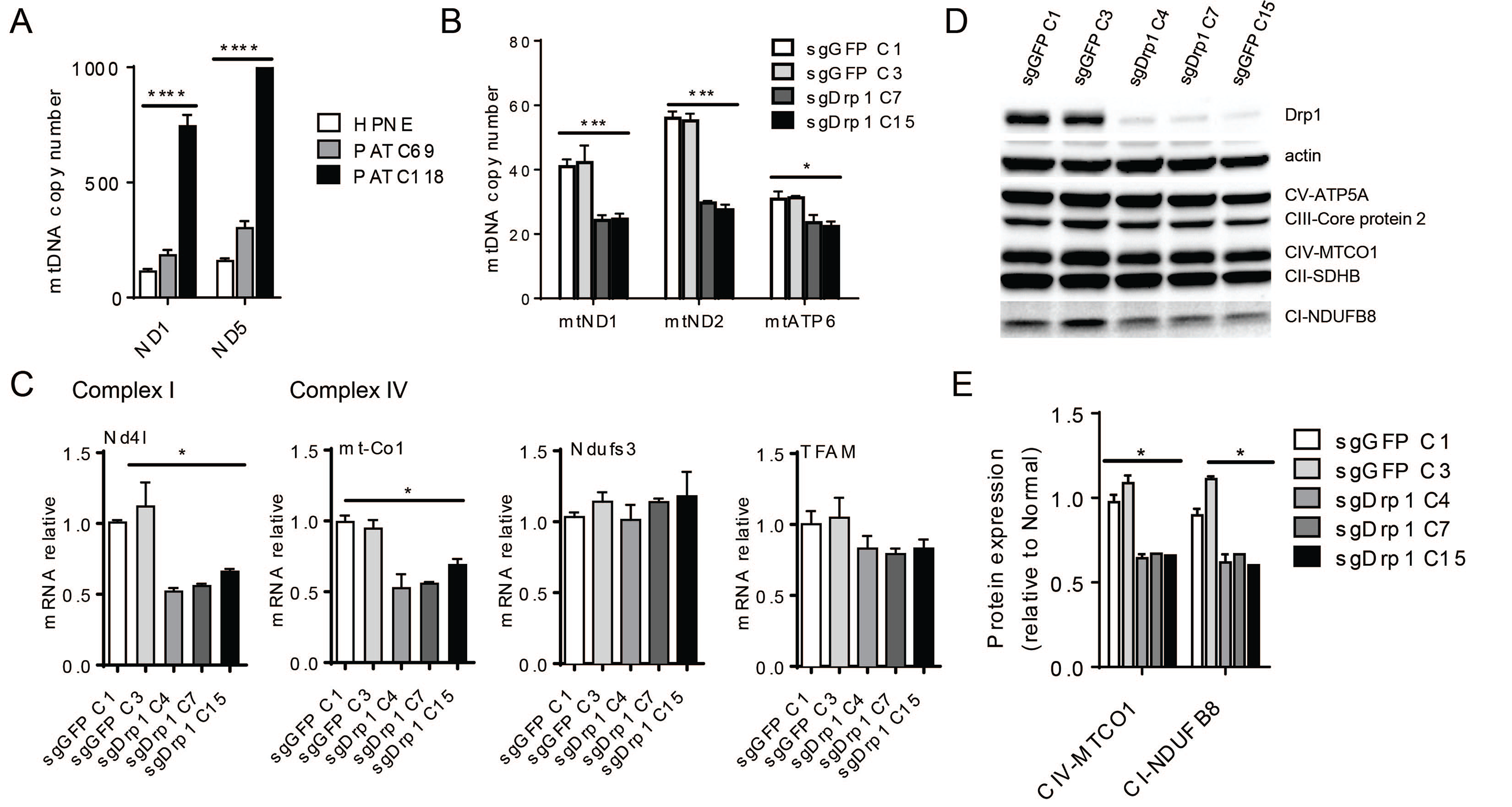
Mitochondrial fusion decreases mtDNA and attenuates mitochondrial Complex I expression. (A) mtDNA copy number of PATC118, PATC69 and human pancreas normal epithelial HPNE. (B) mtDNA copy number of KPC cells with Drp1 knockout. (C) mRNA expression of mtDNA encoded protein and nucleus encoded protein. (D) Western blot of mitochondrial complexes. Including ATP synthase (CV-ATP5 A), succinate-Q oxidoreductase (CII-SDHB), cytochrome-c oxidoreductase (CIII-Core protein 2), NADH dehydrogenase (CI-NDUFB8) and cytochrome C oxidase (CIV-MTCO1). (E) Protein quantification of (D) by measuring band density of complex IV and complex I. Data are presented as mean± SEM. *p<0.05, ***p<0.001, ****p<0.0001.

These mtDNA changes were correlated at mRNA level. Then we detected the mRNA expression of all 13 mitochondrial genes involved in the electron transport chain, and found they were decreased in almost all instances, especially *Nd4l* and *mt-Co1*, both of them were significantly decreased in all three clones compared to control cells (Figure 5C). We measured the expression of nuclear encoded genes involved in the ETC like Ndufs3 and TFAM and found no significant difference, indicating that mitochondrial fusion was likely not inducing global problems with gene transcription.

We then assessed whether these changes at the mtDNA corresponded with alterations at the protein level. Compared to sgGFP controls, protein expression of mitochondrial complex I and complex IV were significantly decreased after Drp1 knockout (Figure 5D, quantified in Figure 5E).

## Discussion

In this study, we demonstrate that inducing mitochondrial fusion suppresses the growth of primary pancreatic cancer lines both in vitro and in vivo by reducing the capacity for oxidative phosphorylation. Mitochondrial function may be critical for cancer progression in the setting of cancer metastasis (LeBleu et al., 2014), and recurrence (Viale et al., 2014), however there are few studies showing its role in primary cancer. To our knowledge, our study is the first to link the inherent metabolic capacity of a cancer cell with mitochondrial dynamics.

Here, we establish that pancreatic cancer metabolism exhibits a dependence on mitochondrial fission. This may be due to the fact that pancreatic cancers nearly uniformly have activating *KRAS* mutations, which then drives fission through Drp1. However, the link between fission and high oxidative phosphorylation has never been previously shown in pancreatic cancer. The mechanisms of the high metabolic drive of pancreatic cancer cells are still not completely clear, but here we demonstrate that it may be at least partially dependent on the dynamics of mitochondria. We show that inhibition of Drp1 or overexpression of the biological antipode Mfn2 induces mitochondrial fusion and reduces oxidative phosphorylation and furthermore suppresses pancreatic cancer growth. Thus, it is likely that the tumor suppression phenotype comes from directly inducing mitochondrial fusion and is less likely a consequence of manipulating Drp1 activity.

We establish a link between mitochondrial fusion and lower oxygen consumption rates and ATP production in the mitochondria and present evidence that this may occur through lower mitochondria mass and reduced expression of Complex I and Complex IV components. The exact mechanism of how mitochondrial fusion leads to decreased mtDNA and ETC activity is unknown, but could be linked through mitophagy (Song et al., 2017) or unfavorable changes in ROS (Balaban et al., 2005) that leads to less functional mitochondria.

Pancreatic cancer survival remains quite low, despite great stride in understanding its biology. Of the major mutations that drive pancreatic cancer, such as *KRAS, TP53, CDKN2A* and *SMAD4*, most of these are not druggable. Here we demonstrate that promoting mitochondrial fusion might a tractable approach to targeting mitochondria in pancreatic cancer. Most normal tissues such as the heart (Eisner et al., 2017), fat, and muscle utilize fused mitochondria for homeostasis, since this helps to repair damaged mtDNA (Chen et al., 2010) and optimizes metabolism (Westermann, 2010). On the other hand, pancreatic cancer exhibits predominantly mitochondrial fission, which is not a feature of most normal mitcohondria. Here, we show that using Mdivi-1 arrests tumor growth and may be an effective treatment. We would predict that the induction of mitochondrial fusion may be highly effective without causing the excessive toxicity that a general mitochondrial inhibitor might have. Future studies should be devoted to finding other small molecules that could induce fusion in other methods (Miret-Casals et al., 2017), which could be tested in clinical trials.

## Material and Methods

### Animal Studies

Animal studies were performed in accordance with the animal protocol procedures approved by the MD Anderson Cancer Center Institutional Review Board and Institutional Animal Care and Use Committee. Orthotopic pancreas injection was performed as shown before (Jiang et al., 2014). C57BL/6J mice were purchased from Jackson laboratories. Tumors were inoculated into the pancreas of recipient C57BL/6 mice with 10^6^ KPC cells in DMEM mixed 1:1 with ice-cold Matrigel (BD Biosciences) for a total of 20 microliters. Mice were imaged twice a week with ultrasound to assess growth. For *in vivo* lung metastasis study, 10^6^ cells were suspend in PBS and injected in mice through tail vein injection. Lungs were fixed and nodules were counted under the dissecting microscope.

### Mdivi-1 Experiments

For *in vivo* Mdivi-1 treatment, mice were heterotopically transplanted with KPC cells in the flank, and when the tumor reached 7mm, mice were assigned to received intraperitoneal vehicle control (10% DMSO) or intraperitoneal Mdivi-1 (10mg/kg) daily for one week. The tumor sizes were measured 2×/week until it reached 15mm, at which time the mice were euthanized and tumors collected for more detailed study.

### Cell culture

Patient derived PATC69, PATC118, PATC50, PATC102 and human normal epithelial HPNE are a kind gift from Dr. Jason Fleming (Kang et al., 2015). Cells were grown in RPMI-1640 supplemented with 2 mM L-glutamine and 10% FBS. PANC198 and murine KPC cells syngeneic with C57BL/6 (K8484) were a generous gift from Dr. Anirban Maitra. The KPC cells were grown in RPMI-1640 with 10%FBS, 2mM GlutaMax, 1mM sodium pyruvate and 7µg/ml insulin. For Tet-On KPC cell lines, the same media except for the substitution of Tet system approved FBS (Clontech). To induce gene overexpression, cells were seeded in 6 well plates in low density and treated with 2µg/ml Doxycyline for 48h. Lenti-X 293T(Clontech) for lentivirals production was maintained in DMEM high glucose with 10%FBS and 2mM GlutaMAX. All cell lines were cultured in a humidified atmosphere containing 5% CO2 at 37°C.

### Plasmids, lentivirals production and clone selection

Human *MFN2* were synthesized and codon-optimized by Genscript then subcloned into PLVX Tet-One puro vector (Clontech). CRISPR gRNA targeting mouse *Dnm1l* was designed at crispr.mit.edu and LentiCRISPRv2 plasmid using standard protocols. The forward sequence is 5’ – CACCGGGTCATGGAGGCGCTGATCC -3’, reverse 5’ – AAACGGATCAGCGCCTCCATGACCC-3’. Lentiviruses were generated using the Lenti-X system from Clontech according to the manufacturer’s instructions.

### Immunoblotting

Cells were lysed in T-PER^TM^ Tissue protein extraction reagent (Thermo Fisher). Cell lysates were resolved on SDS-PAGE gel and transferred onto PVDF membranes using Biorad Trans-Blot Turbo^TM^ transfer system. Primary antibodies were used to identify the relevant protein and loading control (β-actin). Then probed with HRP-conjugated secondary antibodies. The detection of bands was carried out on ChemiDoc imaging system (Biorad). β-actin, DNM1L, mfn2, mfn1 and p-Drp1 616 antibodies are from Cell signaling. Mitochondrial oxidative phosphorylation (OXPHOS) antibody is from Abcam (Esteves et al., 2014).

### Immunofluorescence and Mitochondrial morphology analysis

Cells were seeded in the coverslips with low density, then live cells were loaded with 25nM MitoTracker Red (Thermo Fisher) for 30 minutes, after several washes cells were fixed in 4% paraformaldehyde in growth medium. Coverslips were mounted in mounting medium with DAPI (Zhao et al., 2013); cells were visualized under a confocal microscope (Olympus, FV1000) and processed using Fluoview software (Olympus). Mitochondrial morphology was scored in at least 100 cells per group by using three different categories of mitochondrial morphology (tubular, fragmented and intermediate). Cells with >50% of fragmented mitochondria were scored as mitochondrial fragmentation phenotype. Cells with over 80% of elongated mitochondria were scored as tubular, cells with over 50% short-rod like mitochondria were scored as intermediate (Rolland et al., 2009).

### DNA and mitochondrial DNA copy number

DNA was extracted using DNeasy Blood and Tissue Kit (Qiagen), using 10ng genomic DNA as template for qPCR reaction. Two sets of primers for mtDNA and two sets of primers for nDNA amplification. Nuclear-encoded genes were used as a reference for relative quantification of the mtDNA copy number. For the human cell lines mitochondrial DNA copy number, we use Human Mitochondrial DNA (mtDNA) Monitoring Primer Set (Takara, Cat. #7246). Primers for mtDNA genes are: mtND1 forward 5’-CAG CCG GCC CAT TCG CGT TA-3’, reverse 5’-AGC GGA AGC GTG GAT AGG ATG C-3’; mtND2 forward 5’-TCC TCC TGG CCA TCG TAC TCA ACT-3’, reverse 5’-AGA AGT GGA ATG GGG CGA GGC-3’. The primers for nuclear encoded genes in this study are: nGADPH forward 5’-ACA GCC GCA TCT TCT TGT GCA GTG-3’, reverse 5’-GGC CTT GAC TGT GCC GTT GAA TTT-3’; nNDUFV1 forward 5’-CTT CCC CAC TGG CCT CAA G-3’, reverse 5’-CCA AAA CCC AGT GAT CCA GC-3’.

### Quantitative PCR

Total cellular RNA was extracted using the RNeasy Plus Mini Kit (Qiagen) according to technical specifications. A mix of random primers and oligo (Rolland et al.) were applied for cDNA synthesis using QuantiTech Reverse Transcription kit (Qiagen). Quantitative reverse transcription PCR was performed on a CFX384^TM^ Real-time system (Biorad) using SYBR Green Master mix (Biorad) and gene-specific primers per the manufacturer’s advice. All the PCR primers to amplify mitochondrial encoded proteins are synthesized from IDT (Tan et al., 2015).

### Transmission electron microscopy

Cells were grown and processed in Permanox petri plates as follows: 1X PBS rinse, then fixed at 4°C for 2 days in 2% formaldehyde + 2.5% glutaraldehyde in 0.1M cacodylate buffer, pH7.4 13. After several 0.1M cacodylate buffer rinses, cells were stained in 0.1% tannic acid followed by post-fixation in 1% OsO4 in 0.1M cacodylate buffer + 0.8% potassium ferricyanide. The cells were en bloc stained in 1% aqueous uranyl acetate followed by dehydration through a gradient series of ethanols (50, 70, 80, 90, 95, 100, 100). Cells were then infiltrated with a progressively higher ratio of embedding resin to ethanol, and given 3 changes of pure resin before embedding in the petri plate in a fresh change of Spurr’s Low Viscosity resin 14. Embedded samples were polymerized at 60°C for 3 days then separated manually from the plate, leaving cells on the embedding resin. Ultrathin sections (55-60nm) were cut on a DiaTome Ultra45 knife, using a Leica U7 ultramicrotome. Sections were collected on 150 hex-mesh copper grids and viewed on a Hitachi H7500 transmission electron microscope using an accelerating voltage of 80kV. Images were captured using an AMT XR-16 digital camera and AMT Image Capture, v602.600.51 software.

### Oxygen consumption and glycolytic capacity

Oxygen consumption rate 15 and extracellular acidification rate (ECAR) were determined using the Seahorse XFe24 Extracellular Flux Analyzer (Seahorse Bioscience, North Billerica, MA). Cells were seeded at 20000-40000 per well for measurement. Three electron transport chain (ETC) inhibitors were sequentially injected to each well at specific time points: Oligomycin (2 µM), followed by FCCP (1 µM), followed by the addition of a combination of Rotenone (0.5 µM) and Antimycin (2 µM). For ECAR measurement, cells were plated at the same number as OCR, Three different drugs were sequentially injected to each well at specific time points, here are the final concentration of each drug: Glucose (10mM final concentration, Sigma), oligomycin (2µM final concentration) and 2-deoxy-glucose (100mM final concentration). At the end of each assay, protein quantification was performed for normalization.

### Statistical analysis

Data acquisition and analysis was not blinded. All grouped data are presented as mean ± SEM. Differences between groups were assessed by Student’s t-test or ANOVA using GraphPadInStat software (GraphPad Software, La Jolla, CA).Kaplan-Meier curves were generated and log-rank analysis was performed using GraphPad.

## Acknowledgments

This research was performed in the Flow Cytometry & Cellular Imaging Facility, which is supported in part by the National Institutes of Health through M.D. Anderson’s Cancer Center Support Grant CA016672. TEM imaging for this project was supported by the Integrated Microscopy Core at Baylor College of Medicine with funding from NIH (DK56338, and CA125123), CPRIT (RP150578), the Dan L. Duncan Comprehensive Cancer Center, and the John S. Dunn Gulf Coast Consortium for Chemical Genomics.

## Author contributions

M.Y. and C.M.T. designed the experiments. M.Y. executed most of the experiments. Y.H. created the KPC luciferase cell lines used in in vivo study. T.N.F. and A.D. helped with monitoring tumor growth by ultrasounds and Luminescence. Y.K, E.J.K and J.B.F. created the patient derived pancreatic cancer cell lines used in the study. S.G. and A.M provided the PANC198 and KPC cell line in the study. S.G., J.M.M. and D.L. gave some help with mouse management. The manuscript was written by M.Y. and C.M.T.

